# Direct, sensitive and specific detection of individual single- or double-strand DNA breaks by fluorescence microscopy

**DOI:** 10.1101/772269

**Authors:** Magdalena Kordon, Mirosław Zarębski, Kamil Solarczyk, Hanhui Ma, Thoru Pederson, Jurek Dobrucki

## Abstract

We here describe a technique termed STRIDE (<u>S</u>ensi<u>T</u>ive <u>R</u>ecognition of Individual <u>D</u>NA <u>E</u>nds), which enables highly sensitive, specific, direct *in situ* detection of single- or double-strand DNA breaks (sSTRIDE or dSTRIDE), in nuclei of single cells, using fluorescence microscopy. Sensitivity of STRIDE was tested using specially developed CRISPR/Cas9 DNA damage induction system, capable of inducing small clusters or individual single- or double-strand breaks. STRIDE exhibits significantly higher sensitivity and specificity of detection of DNA breaks than the commonly used TUNEL assay or methods based on monitoring of recruitment of repair proteins or histone modifications at the damage site (e.g. γH2AX). Even individual genome site-specific DNA double-strand cuts induced by CRISPR/Cas9, as well as individual single-strand DNA scissions induced by the nickase version of Cas9, can be detected by STRIDE and precisely localized within the cell nucleus. We further show that STRIDE can detect low-level spontaneous DNA damage, including age-related DNA lesions, DNA breaks induced by several agents (bleomycin, doxorubicin, topotecan, hydrogen peroxide, UV, photosensitized reactions), and fragmentation of DNA in human spermatozoa. STRIDE methods are potentially useful in studies of mechanisms of DNA damage induction and repair in cell lines and primary cultures, including cells with impaired repair mechanisms.

## INTRODUCTION

Decades of studies of mechanisms of DNA damage and repair have led to the development of a number of techniques of detection of various types of DNA lesions. The most sensitive, but indirect and not fully specific^1,2^ techniques of microscopy *in situ* detection of double- or single-strand breaks (DSBs, SSBs) are immunofluorescent staining for phosphorylated histone H2AX (γH2AX)^3^ or recruited repair factors like 53BP1^4^, RAD51^5^ or XRCC1^6,7^. These methods, although relatively sensitive, involve two assumptions: (i) that the repair machinery has been deployed at the site of damage, and that (ii) the DNA lesion is located exactly at the center of the microscopically detectable focus consisting of the recruited repair factors. However, accumulation of repair factors at non-break sites can also occur, thus false positive results are possible^8^. Also, the center of the repair focus may be positioned at a distance from the lesion^9,10^. Direct *in situ* detection of the presence, and determining the spatial position of DNA breaks (i.e. by a chemical reaction with exposed DNA ends) is therefore essential. The two existing techniques that can be used for direct microscopy detection of DNA breaks *in situ, viz.* terminal deoxynucleotidyl transferase dUTP nick-end labeling (TUNEL, for DSBs) or the nick translation (NT, for SSBs) assay^11,12^, usually rely on labeling of accessible DNA ends by procedures which include immunolabeling. Since immunofluorescent detection is limited by typical problems including low signal-to-noise ratio and various levels of nonspecific, and uneven staining^13,14^, sensitivity of these methods does not permit unambiguous detection of the presence and precise location of individual DNA breaks by fluorescence microscopy.

Several new sophisticated, sensitive genome-wide techniques BLESS^15^, BLISS^16^, iBLESS^17^, GUIDE-Seq^18^ and DSBCapture^19^ that can map DSBs to specific genomic loci throughout a cell population have become available in recent years. However, the microscopy toolbox remains very limited^7,11^ and does not offer the matching specificity and sensitivity. Attempts to develop experimental systems to visualize *in situ* single broken DNA ends have been made^20^. These methods, however, enable detection of DSBs only at predetermined sites in the genome.

Here, we present a method abbreviated STRIDE (<u>S</u>ensi<u>T</u>ive <u>R</u>ecognition of Individual <u>D</u>NA <u>E</u>nds), with its two independent variants, which offers unprecedented sensitivity, specificity and ability to reveal precisely the spatial location of single- and double-strand DNA breaks in the nuclei of fixed cells by fluorescence microscopy. This robust tool can detect a DNA break in any nuclear location.

In order to demonstrate an exceptionally high sensitivity of STRIDE we also developed a new, unique methodological approach based on CRISPR/Cas9, which enables simultaneous labeling of a specific genomic locus and induction of one or several closely spaced double-strand cleavages or single-strand nicks at this site in the genome. Cas9 or Cas9n, in combination with truncated gRNA, which were capable of binding to the targeted locus, but had no nuclease or nickase activity were used as labeling control.

## MATERIALS AND METHODS

### Cell culture and cell treatment; sperm cells

HeLa, U2OS cells and human skin fibroblasts were used, and cultured under standard conditions. Human sperms were immobilized on coverslips. Technical details of cell culture and other methods are available in Supplementary Data at NAR online.

### SensiTive Recognition of Individual DNA Ends (STRIDE) dSTRIDE (detection of DSBs)

After cell fixation BrdU was incorporated into DNA ends using terminal deoxynucleotidyl transferase (TdT) (Phoenix Flow Systems, AU: 1001) and detection and fluorescence enhancement was achieved by applying a procedure described in detail in Fig. 1 and Suppl. Materials and Methods (Fig. S1).

**Fig. 1.**
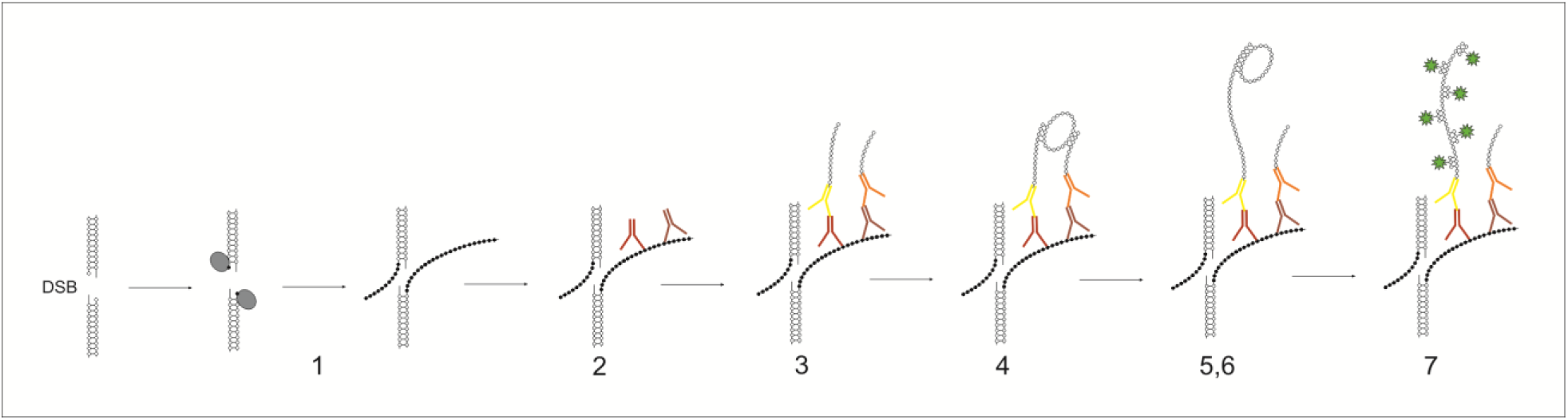
Detecting double-strand DNA breaks by dSTRIDE. Schematic representation of subsequent major steps leading to fluorescent labeling of free DNA ends at the site of a DSB, in fixed cells, by dSTRIDE technique – (1) enzymatic conjugation of nucleotide analogues to DNA ends, (2) attaching primary antibodies of two types (from different hosts), both directed against the incorporated nucleotide analogues, at the concentrations ensuring proximity between the attached antibodies of different types, (3) attaching secondary antibodies with conjugated oligonucleotides to the primary antibodies, (4) hybridizing connector oligonucleotides to two closely located antibody-bound oligonucleotides and ligating them (not shown) to form circular DNA template, (5) rolling circle amplification reaction - oligonucleotides of one of the antibodies acts as a primer for DNA polymerase, (6) synthesizing concatemeric sequences attached to the oligonucleotides on the other antibody by DNA polymerase, (7) hybridizing short fluorescently labeled oligonucleotides to the amplicon (Materials and Methods, and Suppl. Fig. S1 and S2).

### sSTRIDE (detection of SSBs)

Biotinylated and native nucleotides were mixed at a ratio of 3:1 and incorporated at sites of single-strand DNA breaks using *E. coli* DNA Polymerase I. Detection of the incorporated nucleotide analogues and amplification of the fluorescence signals was achieved as described in Suppl. Materials and Methods (Suppl. Fig. S1 and S2).

### Induction of DNA breaks with CRISPR/Cas9 system

In the experimental system we used specific combination of the guide RNAs and the co-expressed fluorescently tagged SpCas9 allowed fluorescent labeling of the targeted repetitive sequence on a long arm of chromosome 3, and simultaneous cutting or nicking DNA in this genomic region. Clusters of DNA cuts or nicks were induced within the repetitive sequence, while individual cuts or nicks were induced at a specific locus immediately adjacent to this repetitive sequence (within distance which is not resolved by standard optical microscopy).

Four types of DNA damage were induced by CRISPR/Cas9 – several closely located nuclease-induced double-strand cuts (which we call ‘clusters of cuts’ for brevity), individual cuts, clusters of nickase-induced nicks, and individual nicks.

In order to induce clusters of double-strand cuts CRISPR/Cas9 nuclease with appropriate guide RNAs (which decreased the nuclease activity of Cas9), targeted to a specific repetitive sequence in a long arm of chromosome 3, was used. A cluster of DSBs was thus induced. At the same time DNA-bound SpCas9-3XGFP served as a fluorescent label for this genomic locus.

Individual cut was induced by targeting enzymatically active Cas9-gRNA complex to a selected locus near the repetitive sequence on chromosome 3, as described in detail in Suppl. Materials and Methods. In the case of individual cut fluorescent labeling of the adjacent repetitive sequence was achieved by targeting fluorescently tagged Cas9 to this locus on chromosome 3 by truncated gRNA, which yielded the DNA-bound Cas9-gRNA complex inactive.

In order to induce clusters of nicks (single-strand DNA breaks) CRISPR/Cas9n (a mutated Cas9 protein, exhibiting nickase activity) with appropriate guide RNA, targeted to the abovementioned specific repetitive sequence in a long arm of chromosome 3, was used. A number of SSBs were induced in this chromosome region. As in the case of clusters of cuts, also in the case of nicks DNA-bound SpCas9n-3XGFP served as a fluorescent label for this genomic locus.

Individual nick, accompanied by labeling of the adjacent chromosomal region, was induced by targeting enzymatically active Cas9n-gRNA complex to a selected genomic locus near the repetitive sequence on chromosome 3, as described in detail in Suppl. Materials and Methods. Fluorescent labeling of the adjacent repetitive sequence was achieved by targeting SpCas9n-3XGFP, using short gRNA.

Plasmid construction, transfection procedures, detailed description of this experimental system and other relevant information on methods is available in Supplementary Data at NAR online.

## RESULTS

### STRIDE principle

The detection of endogenous or induced single- or double-strand DNA breaks by STRIDE consists of three major steps: (i) conjugation of deoxynucleotide analogues to exposed free DNA ends (3’-OH) by terminal deoxynucleotidyl transferase (TdT) or DNA polymerase I (Pol I) (step 1, Fig. 1, and Suppl. Fig. S1), (ii) specific recognition of the incorporated nucleotides at DNA ends by a mixture of two (or more if appropriate) different antibodies (at carefully optimized concentrations resulting in antibodies of both types bound side-by-side at the target) against this target molecule (steps 2-3, Fig. 1, and Suppl. Fig. S1), and finally (iii) fluorescence signal enhancement based on rolling circle amplification reaction (RCA) and detection of the amplified DNA by hybridization with fluorescently labeled oligonucleotides (steps 4-7, Fig. 1 and Suppl. Fig. S1). The combination of direct reaction with DNA ends, binding of antibodies to the conjugated nucleotide analogues, and rolling circle amplification reaction (occurring only with two adjacent antibodies of different types located close to each other) followed by hybridization of numerous fluorescent probes results in strong signal amplification and near-zero signal background in microscopy images where even individual DNA breaks are represented by bright, readily detectable fluorescent signals (Suppl. Fig. S2). The critical point of the method is the use of two antibodies raised in different hosts, directed against a chain of identical components, yet avoiding the use of fluorescently labeled secondary antibodies – this particular approach makes it possible to take advantage of high signal amplification offered by proximity ligation assay, and avoiding typical background signal always associated with the detectable nonspecific binding of the antibodies in immuno-fluorescence assays. Very low or undetectable background and a lack of nonspecific signals are preconditions for high sensitivity detection of individual molecular events like DNA breaks – this condition is fully satisfied by STRIDE (for further discussion of the pertinent technical issues and variants of the technique see Materials and Methods, Suppl. Fig. S1, and Suppl. Fig. S2).

### Sensitivity of STRIDE – detection of Cas9-induced DNA cuts and nicks

To test sensitivity of a new method it is first necessary to induce a known and controllable low number of DNA breaks in the nucleus to be detected. DNA lesions occurring all throughout the nucleus can be induced by a number of methods, including exposure of the whole cell to UV, ionizing radiation, or certain DNA-damaging drugs. In these cases spatial positions of the induced DNA lesions cannot be predicted, nor can such a damage be distinguished from endogenous DNA breaks. DNA damage can be induced in a defined nuclear region by exposing a selected site within the nucleus to a focused beam of visible light, in the presence or absence of DNA-bound photosensitizers (microirradiation^6,21,22^). In all these approaches the type of DNA damage (oxidative, DNA breaks of various types), the number and the exact spatial or genomic positions of the induced lesions is not under full control of an experimenter. Thus, we developed CRISPR/Cas9-based experimental system capable of inducing well defined damage (single- or double-strand breaks; no base modifications or oxidative damage) in selected loci in the genome. Using specially adapted CRISPR/Cas9 we *in situ* induced small clusters or individual, double-strand or single-strand DNA breaks resulting from the cleavage activity of nuclease SpCas9, or the nicking activity of the non-target strand-cleaving nickase version of SpCas9 with HNH domain deactivated by introduction of a point mutation H840A^23,24^ We used engineered guideRNAs with nuclease-active SpCas9 tagged with 3XGFP or 3XmCherry, or nickase-active SpCas9n tagged with 3XGFP^25^. They were employed to induce DNA lesions and simultaneously visually detect the sites of their induction inside cell nuclei. We targeted a subtelomeric region of the long arm of chromosome 3 which contains a repetitive sequence that is unique to this site and this chromosome^26^. Subsequently we detected the Cas9-induced DNA damage by sSTRIDE or dSTRIDE.

Clusters of double-strand breaks induced by CRISPR/Cas9 were readily detected by dSTRIDE (Fig. 2a-c). The foci representing the target-bound Cas9 itself (Fig. 2a) and the dSTRIDE signal (Fig. 2b) colocalize (Fig. 2c, seen also in magnified views and profiles of fluorescence signals in panels c-1 and c-2) confirming that Cas9-induced DSBs were detected by dSTRIDE. The expected accumulation of a repair factor 53BP1, and phosphorylation of histone H2AX, were also observed at the sites of accumulation of SpCas9 (data not shown). In parallel we tested the capacity of TUNEL, the only available method of direct *in situ* detection of DSBs, to detect Cas9-induced DSBs. Fig. 2, panels a-c vs. d-f, show a comparison between fluorescence signals representing clusters of DSBs detected by dSTRIDE and TUNEL in nuclei of cells in which the subtelomeric repeats were cleaved by Cas9. Distinct dSTRIDE foci were readily observed (Fig. 2b-c) whereas no signal was detected with TUNEL (Fig. 2e-f), despite the fact the SpCas9 was clearly present at the targets in both cases (Fig. 2a and d, presented also in magnified views and profiles of fluorescence signals in panels f-1 and f-2). We also used dSTRIDE and TUNEL to detect DNA ends generated by DNase I (Fig. 3) and compared sensitivity of both methods (Fig. 3d and h). We conclude that TUNEL did not have the sensitivity required to either detect clusters of closely localized DNA breaks induced by Cas9 or the low level damage induced by DNase I.

**Fig. 2.**
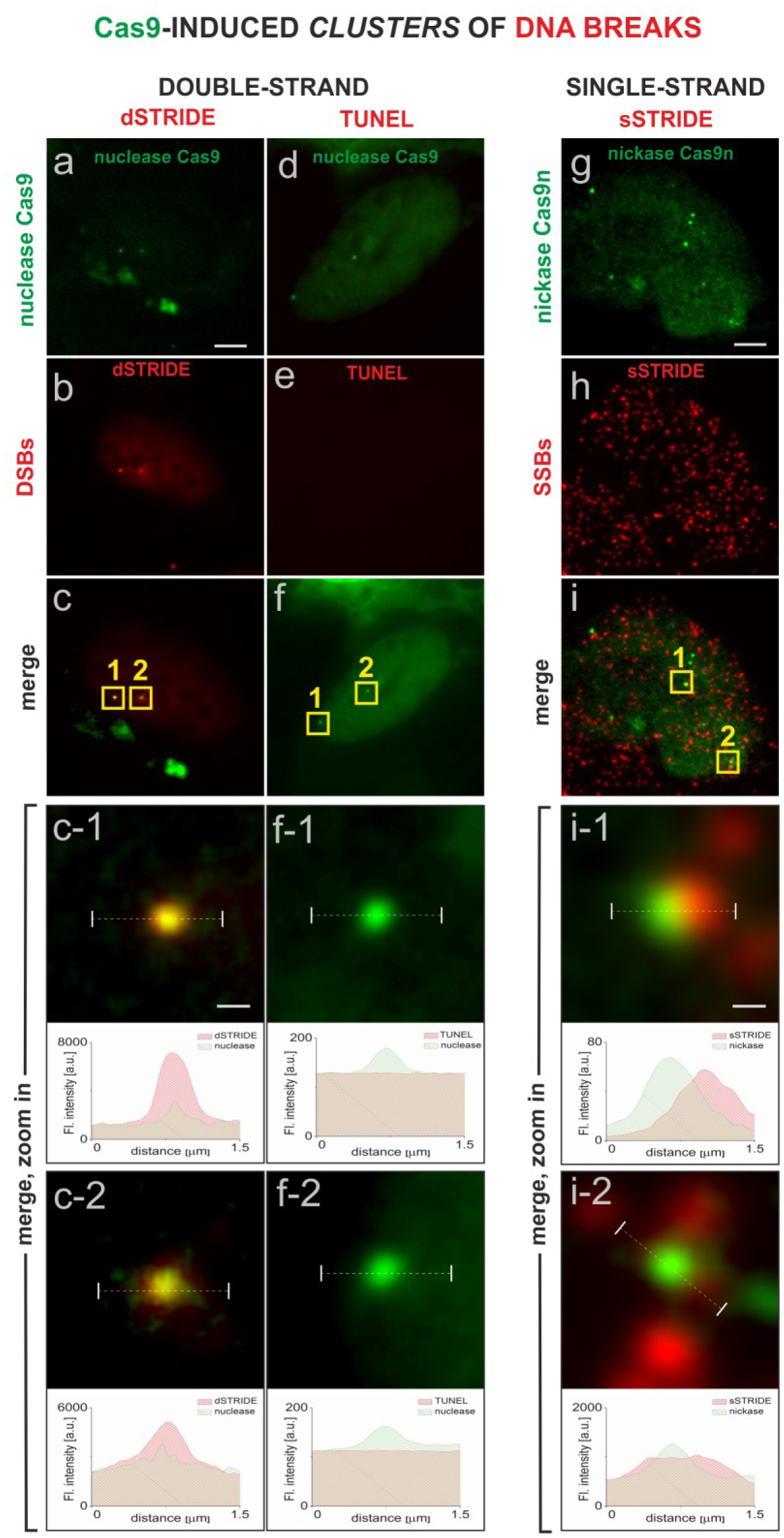
Imaging clusters of double- or single-strand DNA breaks by STRIDE. DNA lesions were induced by SpCas9 in a unique region of chromosome 3 that contains chromosome-specific repetitive sequence (Suppl. Fig. S5). The genomic loci targeted by SpCas9 are detected as green fluorescent foci due to the presence of many chromatin-bound 3XGFP-labelled SpCas9 molecules^25^. Distinct fluorescent dSTRIDE or sSTRIDE foci colocalize with the sites of SpCas9 or SpCas9n accumulation, confirming an ability of STRIDE to detect clusters of DNA lesions localized on a long arm of chromosome 3. TUNEL assay does not have sensitivity to detect SpCas9–induced DNA lesions. Scale bars: 5 µm and 0.3 µm (zoom in). **(a-c)** detecting DNA-bound Cas9, and clusters of Cas9 nuclease-induced double-strand breaks: **(a)** green foci representing SpCas9-3XGFP nuclease accumulated in genomic loci on chromosome 3, **(b)** red foci of dSTRIDE signal, associated with clusters of double-strand DNA breaks, **(c)** merge of SpCas9 nuclease and dSTRIDE signals demonstrating an ability to detect the presence and spatial positions of DNA breaks in the region of accumulation of SpCas9 on chromosome 3, **(c-1, c-2)** magnified views of the overlapping foci of SpCas9 and dSTRIDE (yellow squares in panel c) at the two intranuclear locations, and the corresponding fluorescence profiles showing relative positions of these foci, **(d-f)** clusters of double-strand DNA breaks induced in the two CRISPR-targeted loci by SpCas9-3XGFP (green) are not detected by TUNEL assay (red): **(d)** green foci representing SpCas9-3XGFP accumulated in genomic loci on chromosome 3, **(e)** weak red background TUNEL signal present throughout the nucleus, but no red foci or areas of higher fluorescence intensity can be detected at the sites of SpCas9 accumulation, **(f)** merge of SpCas9-3XGFP and TUNEL signals, **(f-1, f-2)** magnified views of the two SpCas9 nuclease accumulation foci (panel f, yellow squares), showing no TUNEL signal at the sites of DNA lesions induced by SpCas9, and the corresponding fluorescence profiles showing the positions of Cas9 foci and an absence of TUNEL signal at these sites. **(g-i)** detecting DNA-bound Cas9n, and clusters of Cas9n nickase-induced single-strand breaks: **(g)** green foci representing SpCas9n-3XGFP nickase accumulated at genomic loci on chromosome 3, **(h)** numerous red foci of sSTRIDE signal, associated with numerous endogenous, and SpCas9n-induced, single-strand DNA breaks, **(i)** merge of SpCas9n (in the region of accumulation of SpCas9n on chromosome 3), and sSTRIDE signals (representing DNA ends), demonstrating an ability to identify SpCas9n-induced single-strand DNA breaks among many endogenous SSBs, **(i-1, i-2)** magnified views of the overlapping foci of SpCas9n and sSTRIDE (yellow squares in panel i) at the intranuclear locations, where SSBs were induced by Cas9n. Corresponding fluorescence profiles show relative positions of these foci.

**Fig. 3.**
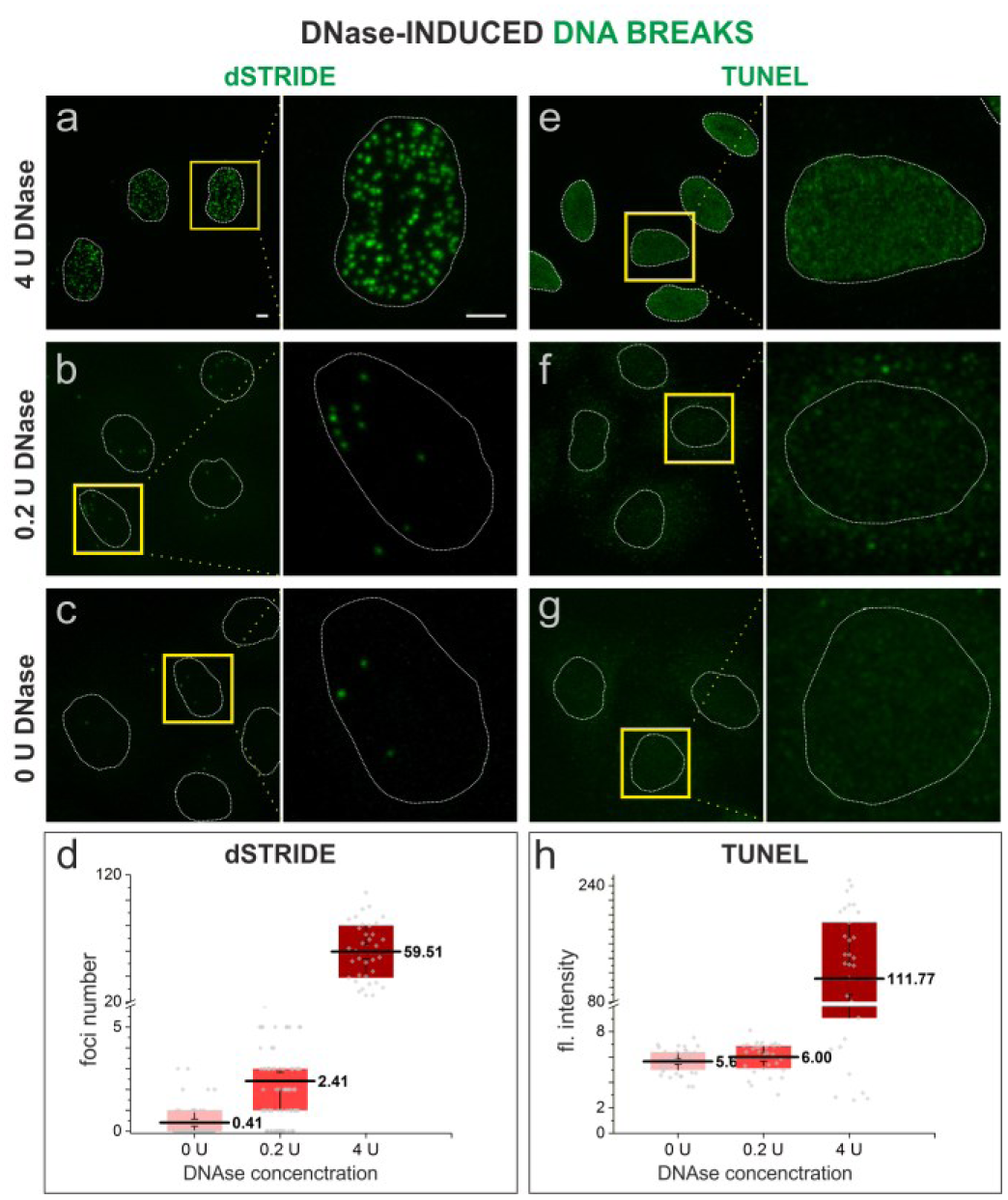
dSTRIDE is more sensitive than TUNEL. To validate the dSTRIDE method and compare the sensitivity of dSTRIDE (a - d) with TUNEL (e - h) free DNA ends were induced in fixed HeLa cells by exposure to DNase at low (0.2 U) or high (4 U) concentration, and detected using either method. For both techniques, treatment of cells with DNase I, leading to induction of numerous DNA ends, resulted in significant fluorescence signals in the treated nuclei when compared to untreated cells (c, g - where only DNA breaks resulting from low level endogenous damage are to be expected). **(a-c, e-g)** Representative fluorescence confocal images (and magnified views) of untreated and DNase I-treated HeLa cells in which DNA ends were detected by dSTRIDE (a-c) or TUNEL (e-g) are shown. For both assays, treatment of cells with a high concentration of DNase I (4 U) (a, e) resulted in a higher signal in the nuclei than in untreated cells (0 U) (c, g). However, only for dSTRIDE a difference between cells treated with a low concentration of DNase I (0.2 U) (b) and untreated cells (0 U; only endogenous damage) (c) was detectable – the number of detected DSBs was significantly higher in DNase treated cells. For TUNEL assay, no significant fluorescent signal was observed in samples treated with a low concentration of DNase I (0.2 U; low number of induced DNA ends) (f) when compared to untreated cells (0U, i.e. exclusively endogenous damage) (g). Scale bars – 5 µm. **(d, h)** Box plots representing the results of analysis of microscopic images of DNA breaks. For TUNEL (h) assay the signal, due to its characteristic blurriness, was the mean grey value of pixels (fluorescence intensity) within the nucleus (note that individual lesions or groups of lesions were not detectable). For dSTRIDE (d), the number of foci (representing individual lesions or their clusters) in each nucleus was determined. The bottom of each box is the 25th percentile, the top is the 75th percentile. The solid horizontal line represents the mean value, the whiskers represent standard error of the mean (s.e.m.). An independent two-sample t-test has shown that the difference in mean values between samples treated with a low concentration of DNase I (0.2 U; low number of induced DNA ends) and untreated samples (0 U; only endogenous DNA damage) was statistically significant only in dSTRIDE (d) (p-value=2.8 × 10^−7^).

We then investigated whether the clustered DNA nicks produced by nickase SpCas9n can be detected by sSTRIDE (Fig. 2g-i). In this case sSTRIDE was also shown to be very sensitive. SSBs were readily detected by sSTRIDE (XRCC1, a repair factor involved in SSB repair was recruited to Cas9n-induced damage, as expected; data not shown). Fig. 2h shows a large number of sSTRIDE signals marking single-strand DNA breaks, reflecting the known existence of numerous endogenous SSBs in cell nuclei. As anticipated, colocalization of sSTRIDE signals with SpCas9n was also observed, demonstrating that the clusters of SSBs induced by Cas9n were readily detected by sSTRIDE (Fig. 2g-i).

STRIDE was capable of detecting not only clusters, but even individual double- or single-strand DNA breaks induced at the sites of CRISPR/Cas9 accumulation (Fig. 4a-f). As expected, in a control experiment, when inactive complexes of nuclease Cas9 or nickase Cas9n with short guide RNAs were used resulting in no cleavages, there was no colocalization between dSTRIDE and Cas9, or sSTRIDE and Cas9n foci (not shown, in this case STRIDE signals represented only endogenous DNA breaks, see also Suppl. Fig. S3). When an active Cas9/gRNA or Cas9n/gRNA complex was used, an individual DNA cut or nick was induced and detected by STRIDE. Fig. 4 shows CRISPR/Cas9-induced individual DSBs or SSBs detected by dSTRIDE (Fig. 4a-c; see also quantitative analysis of the number of foci in Suppl. Fig. S3a) and sSTRIDE (Fig. 4d-f,Suppl. Fig. S3b), respectively. To the best of our knowledge, the images in Fig. 4 are the first examples of direct fluorescence-based microscopy *in situ* detection of individual DNA ends induced by CRISPR/Cas9.

**Fig. 4.**
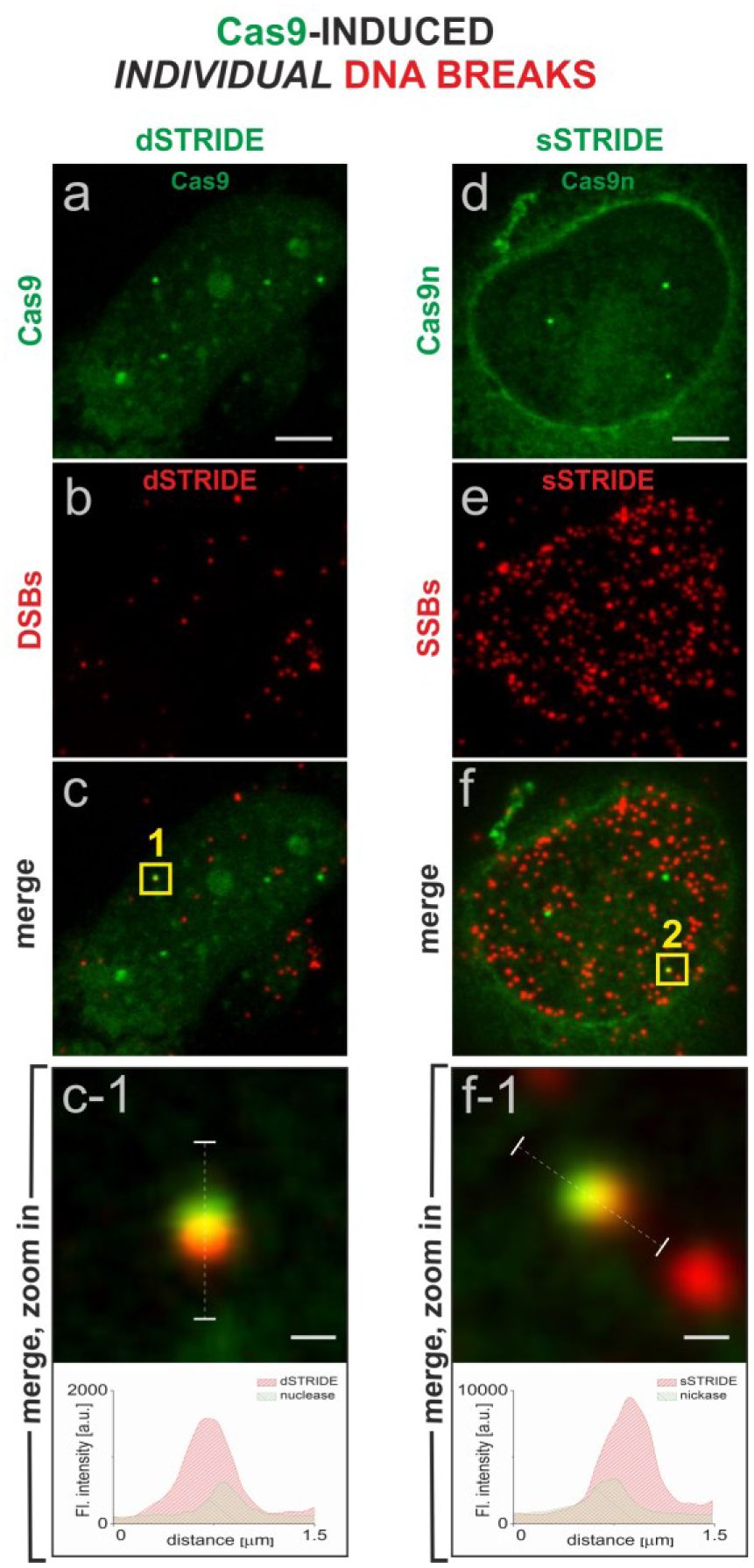
High sensitivity of STRIDE demonstrated by imaging individual single- or double-strand DNA breaks. The individual DSBs and SSBs were induced by targeting a fluorescently labelled nuclease (SpCas9, green) (a) or nickase (SpCas9n, green) (d) to a unique sequence on chromosome 3 (Suppl. Fig. S5; details in Materials and Methods). Scale bars: 5 µm and 0.3 µm (zoom in). sSTRIDE and dSTRIDE are both capable of detecting *individual* CRISPR/SpCas9- and CRISPR/SpCas9n-induced double- or single-strand DNA breaks, as shown by colocalization of the green (Cas9) and red (STRIDE) foci (c, f), yielding yellow areas. **(a-c)** individual DSB detected by dSTRIDE (a - SpCas9, green; b – dSTRIDE, red; c – merge of both signals; c-1 - a magnified view of the damage site, yellow square at panel c) **(d-f)** individual SSB detected by sSTRIDE (d - SpCas9n, green; e – sSTRIDE, red; f – merge of both signals; f-1 - a magnified view of the damage site, yellow square at panel f)

### Detecting endogenous and induced DNA breaks

The mechanisms of damage induction and the type of resulting damage vary substantially between different damaging stimuli. Thus, we tested an ability of STRIDE to detect DNA breaks arising throughout the nucleus from the action of various agents, in cells of various types and chromatin of different levels of condensation.

As shown in Fig. 5, STRIDE was found to be capable of assessing low levels of endogenous double-strand DNA breaks in fibroblasts obtained from 3 and 60 year old donors and maintained in primary cell cultures (Fig. 5a and b). Quantitative analyses of the number of detected double-strand breaks showed that cells from a 60 year old subject contained significantly more double-strand breaks (Fig. 5b and c), as would be expected based on the existing indirect evidence^27^. Moreover, STRIDE was also capable to demonstrate and measure differences between the numbers of double-strand DNA breaks induced by antitumor anthracycline antibiotic doxorubicin in cells from a young and elderly donor, and demonstrated that susceptibility to drug-induced damage was significantly higher in the cells from a 60 years old donor (Fig. 5c). Interestingly, it also showed that the numbers of DNA breaks induced with doxorubicin varied more widely between cells from an older donor.

**Fig. 5.**
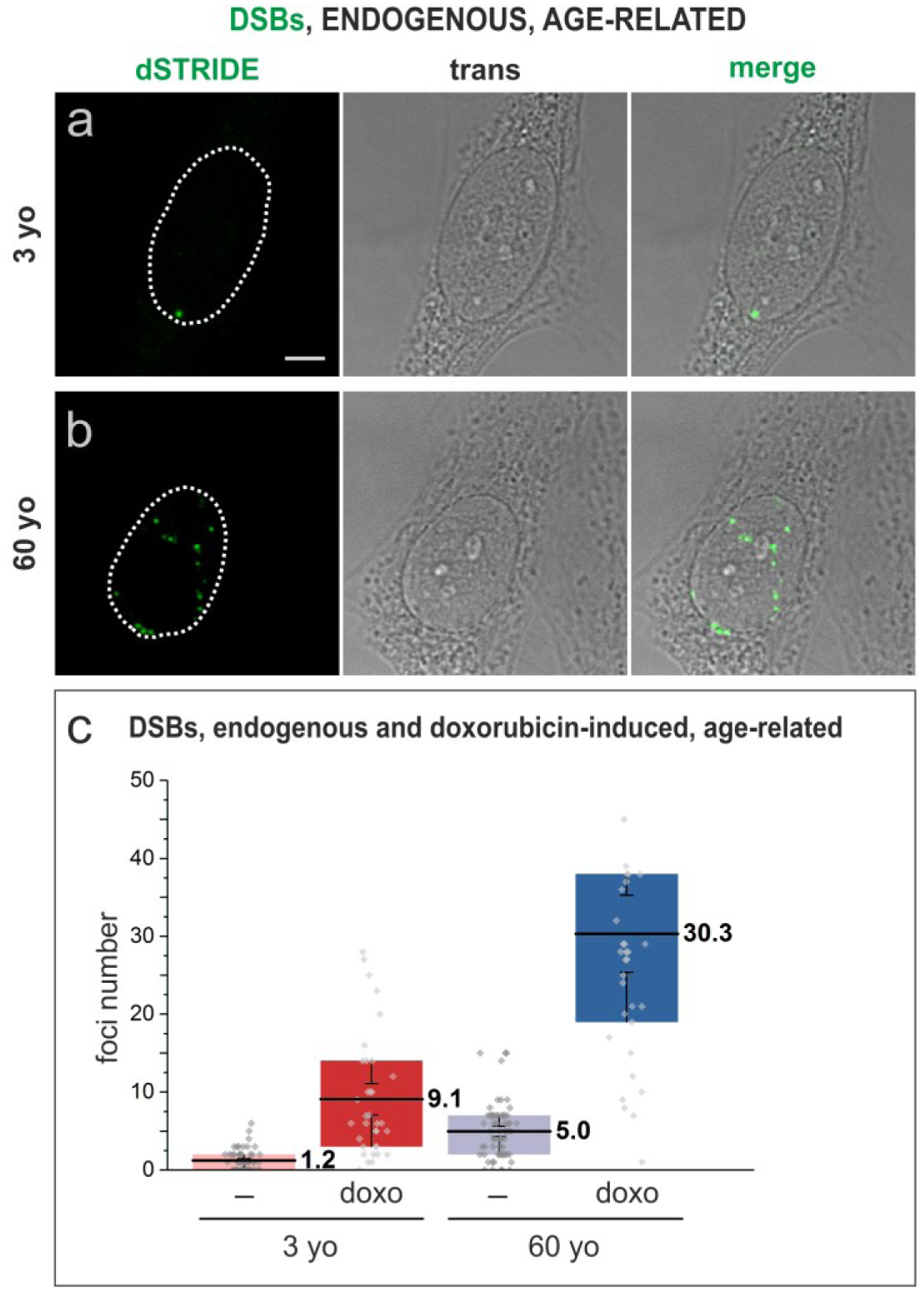
Imaging DNA breaks, endogenous or experimentally induced by doxorubicin, in human fibroblasts from a 3 or 60 year old donors in primary cultures. **(a**,**b)** fluorescence signals marking double-strand breaks detected by dSTRIDE (green) in fibroblasts (transmitted light images) from 3 (a) or 60 year old (b) donors. Scale bar: 5 µm. **(c)** the numbers of endogenous and doxorubicin-induced double-strand DNA breaks detected in fibroblasts from a 3 year old and 60 year old donor, demonstrating a higher number and a larger range of numbers of endogenous DNA lesions in cells from an older donor, and showing a higher number of lesions induced by the drug in cells of an elderly individual. Box plots represent the results of analysis of microscopic images of DNA breaks in which the number of foci (representing individual lesions or their clusters) in each nucleus was determined. The bottom of each box is the 25th percentile, the top is the 75th percentile. The solid horizontal line represents the mean value, the whiskers represent standard error of the mean (s.e.m.).

We subsequently tested an ability of STRIDE to detect DNA breaks induced by various conditions and damaging treatments that are known to result in various levels and types of DNA damage. DNA breaks in cells undergoing spontaneous apoptosis showed very high levels of dSTRIDE signal, as expected (Supp. Fig. S4). DNA breaks induced in cells of an established cell line by radiomimetic antitumor antibiotic bleomycin (Fig. 6), chemical factors including hydrogen peroxide (Fig. 7a), topoisomerase inhibitor topotecan (Fig. 7b), and by exposure to UV (Fig. 7c) were also readily detected by dSTRIDE, regardless of the nature of the factor inducing damage.

**Fig. 6.**
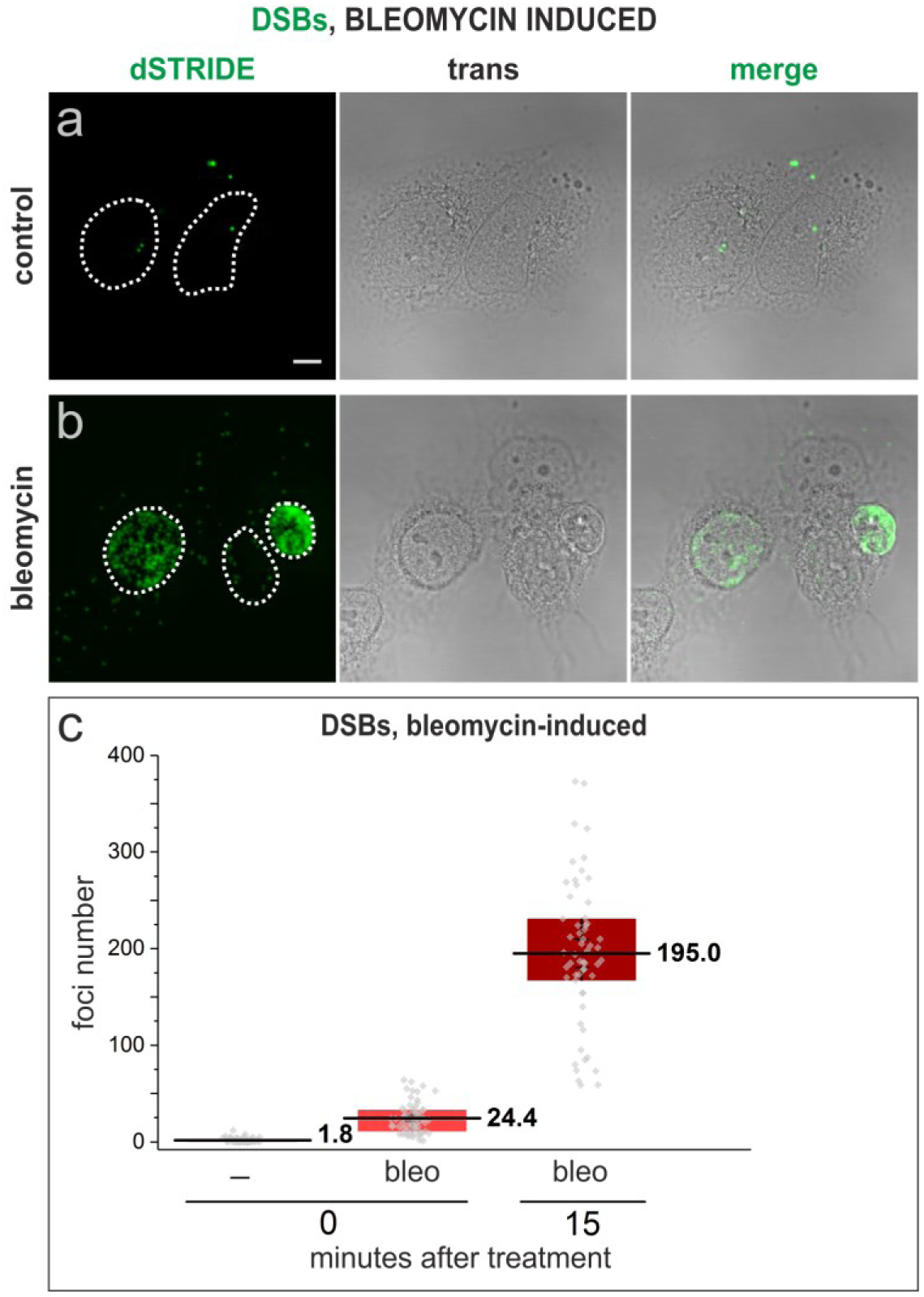
Imaging DNA breaks induced in U2OS cells in an *in vitro* culture, following exposure to a radiomimetic antitumor antibiotic bleomycin. **(a, b**) control cells, showing only low level endogenous DNA damage (a) and cells treated with bleomycin (b) that show signs of heavy damage or apoptosis (the cell nucleus presented on the right-hand side of the image) (dSTRIDE fluorescence signals, transmitted light image showing cell morphology, and an overlay of both images). Scale bar: 5 µm. **(c)** quantitative analysis of the numbers of DSBs in control cells, and in cells exposed to bleomycin for 30 min. and fixed and stained for STRIDE immediately (labeled with 0) or 15 min. (labeled with 15) after exposure to bleomycin, showing an increase of the number of DNA lesions during 15 min. after exposure to the drug. Box plots represent the results of analysis of microscopic images of DNA breaks in which the number of foci (representing individual lesions or their clusters) in each nucleus was determined. The bottom of each box is the 25th percentile, the top is the 75th percentile. The solid horizontal line represents the mean value, the whiskers represent standard error of the mean (s.e.m.).

**Fig. 7.**
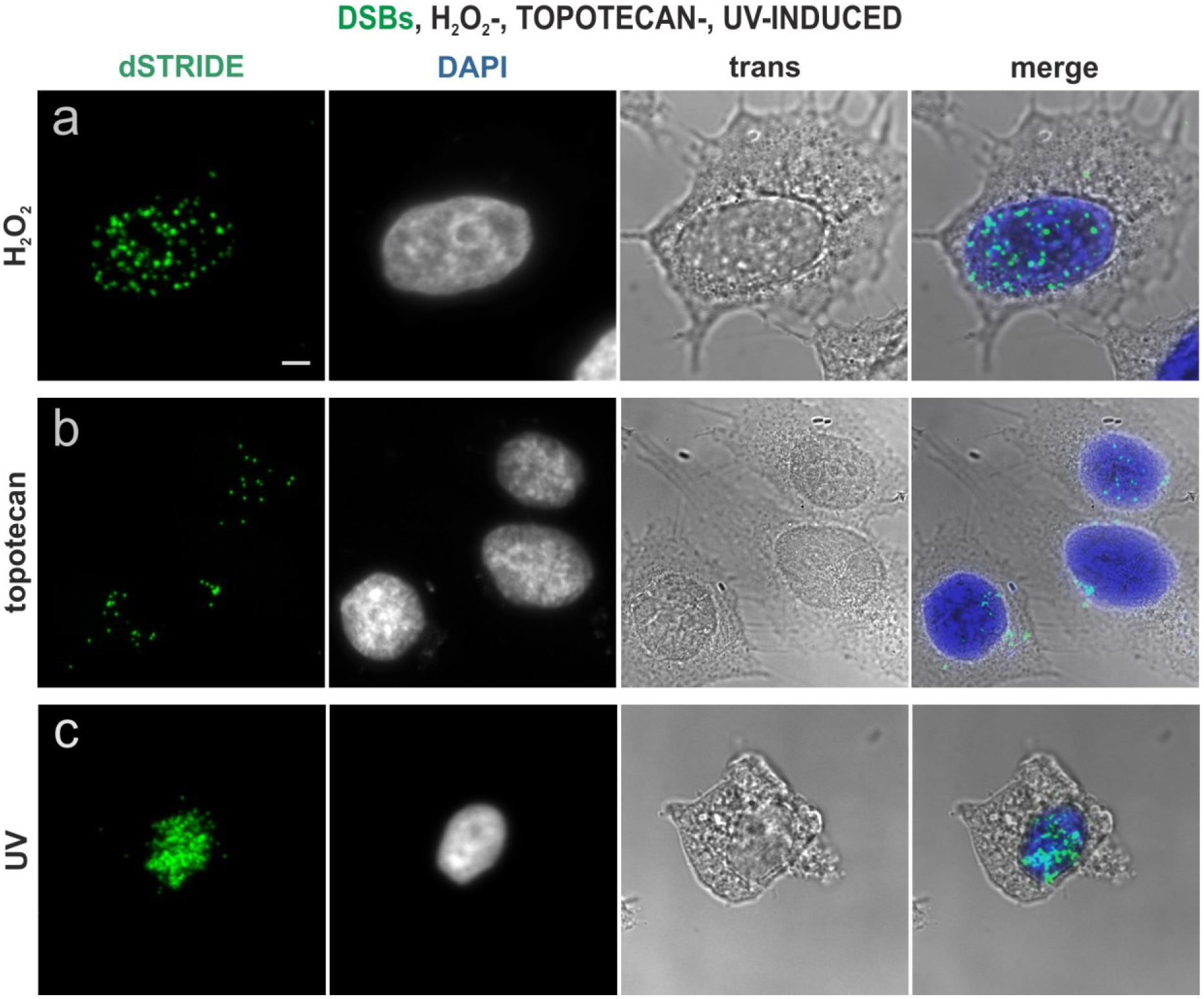
Imaging DNA breaks induced in HeLa cells in an *in vitro* culture, by various damaging agents. Double-strand DNA breaks induced by exposure to hydrogen peroxide (a), topoisomerase inhibitor topotecan (b), and ultraviolet light (c), and detected by dSTRIDE are shown. STRIDE signals (green), DAPI-stained nuclei, transmitted light images and merged images (with DAPI in blue) are shown. Scale bar: 5 µm.

dSTRIDE also detected DNA breaks induced by laser microirradiation in a selected region of the nucleus. This approach is often used in order to image recruitment of repair factors to the site of damage. When a focused light beam was applied to a small region of the cell nucleus, in the presence of a DNA-bound photosensitiser^21^ (ethidium bromide) (Fig. 8), oxidative damage and DNA breaks were induced^21^. As shown in Fig. 8a-b induction of DSBs in a selected area in the cell nucleus was confirmed by dSTRIDE. TUNEL, as previously noted, was not capable of detecting these lesions (Fig. 8c-d).

**Fig. 8.**
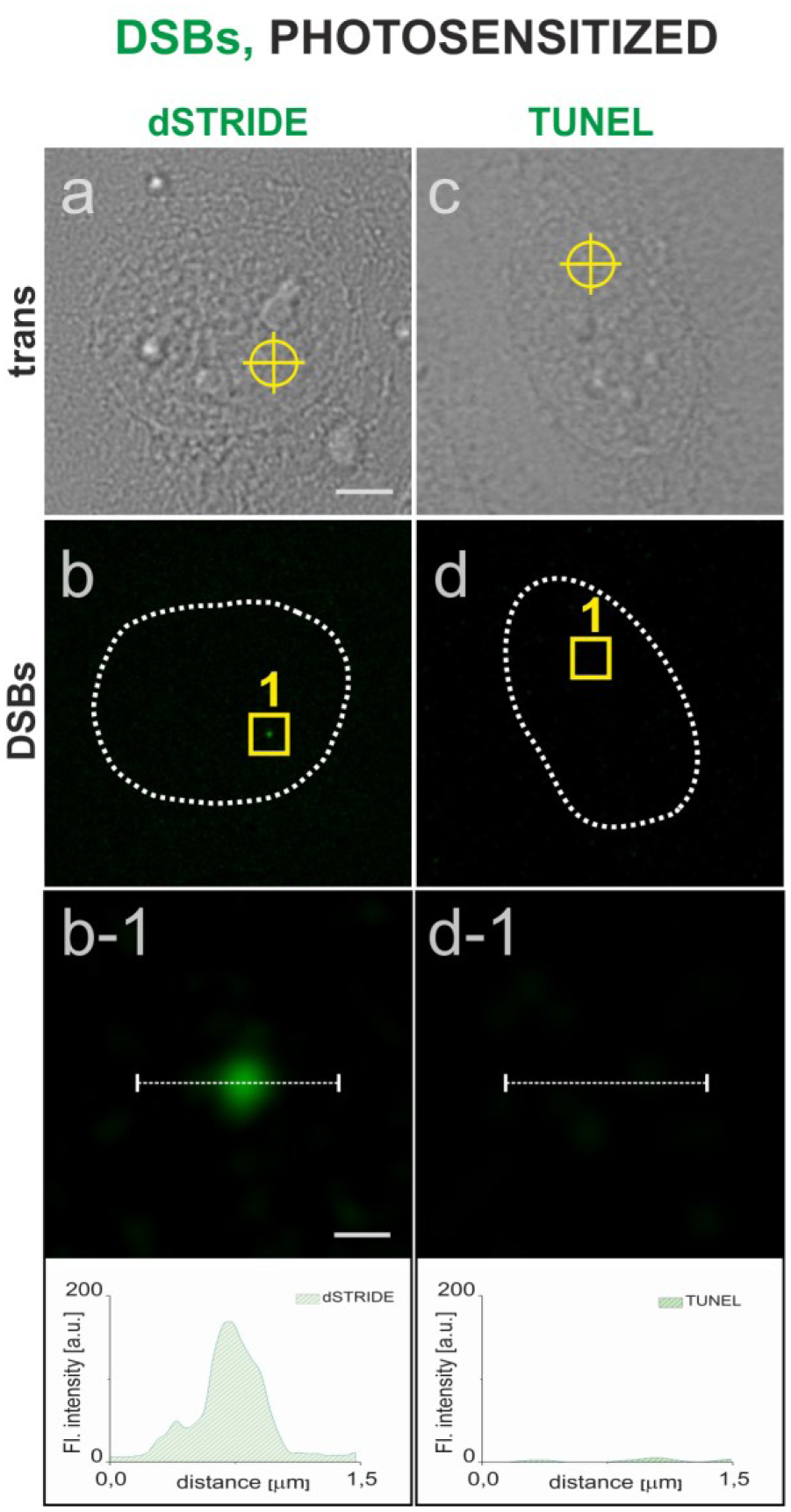
Detection, by dSTRIDE and TUNEL assay, of DNA breaks inflicted in a small selected region of the cell nucleus by photosensitized reactions. DNA-bound ethidium anion was used as a photosensitizer (500 nM ethidium bromide in culture medium)^21^; ethidium was excited by local illumination with 488 nm focused laser beam (300 nm diameter). Scale bars: 5 µm and 0.3 µm (zoom in) **(a-b)** transmitted light image of a cell, with a site at which the laser beam was focused marked by a circle (a), an image showing a fluorescence dSTRIDE signal at the site of damage (b), a magnified view of the region with the damage site (b-1), and a corresponding fluorescence profile, demonstrating that dSTRIDE detected the local photosensitized damage; **(c-d)** transmitted light image of a cell showing the site of induction of photosensitized damage (c), fluorescence image (d), and a magnified view as well as the corresponding fluorescence profile (d-1) demonstrating a lack of TUNEL signal at the site of photosensitized damage.

Finally, in order to test versatility of STRIDE we examined an ability of this method to detect DNA damage in spermatozoa, lymphocytes and tissue sections. DNA fragmentation is an important parameter which characterizes the quality of sperm cells and its level is frequently tested in male infertility assessment. We attempted to detect single-strand breaks and DNA fragmentation in human sperm cells of a patient of a fertility clinic, using sSTRIDE and dSTRIDE. When multistep labeling procedure is used, spermatozoa pose different technical problems than adherent cells due to a lack of attachment to substratum, fragility and highly condensed state of chromatin. However, when a procedure of maintaining sperms immobilized on glass support was optimized (Materials and Methods), DSBs and SSBs were detected successfully by dSTRIDE and sSTRIDE (Fig. 9a,b). Endogenous DNA breaks were also successfully detected in lymphocytes and Formalin-Fixed Paraffin-Embedded as well as frozen mouse tissue sections (data not shown).

**Fig. 9.**
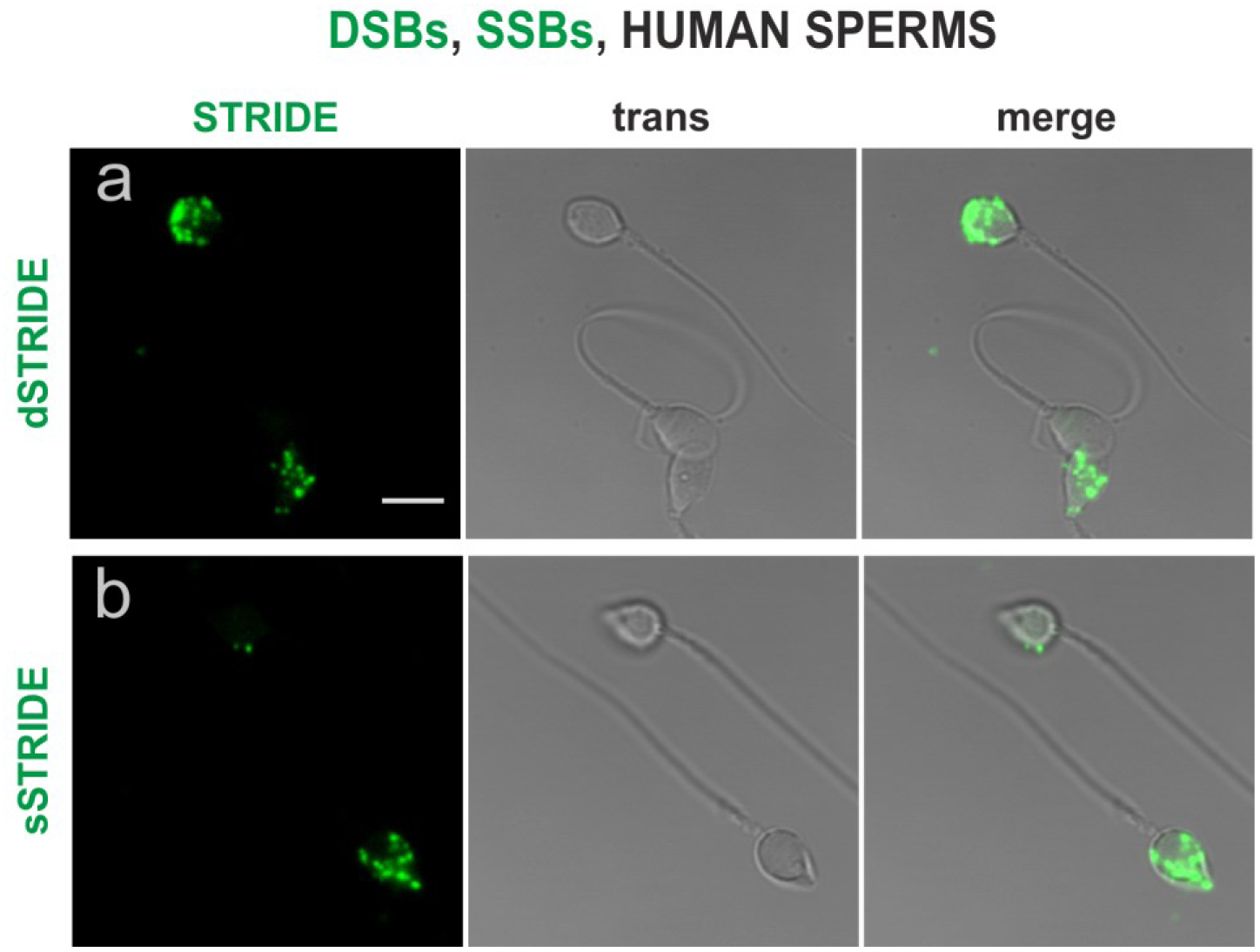
DNA fragmentation in spermatozoa - detecting endogenous levels of DNA damage (single- and double-strand breaks) in human sperm cells by dSTRIDE and sSTRIDE. **(a, b)** Examples of low and high levels of DNA double-strand (a) and single-strand (b) breaks in sperms of a fertility clinic patient are shown (dSTRIDE fluorescence signals, transmitted light, and merged images). Scale bar: 5 µm.

## DISCUSSION

The data presented demonstrate that dSTRIDE and sSTRIDE are capable of direct *in situ* detection of small clusters and even individual single- and double-strand DNA breaks, whether endogenous or induced experimentally in cellular chromatin by factors and treatments of various types. STRIDE methods rely on labeling procedure which maintains the nonspecific signal at an almost undetectable level, and on strong amplification of damage-specific fluorescence signal. STRIDE methods are versatile in that DNA breaks resulting from treatments with various DNA damaging agents and exposures to various damage conditions, in cultured cells lines, cells from patients maintained in primary cultures, human spermatozoa, lymphocytes and tissue sections can be readily detected and imaged by standard fluorescence confocal microscopy.

Due to exceptional sensitivity both dSTRIDE and sSTRIDE have considerable potential to become powerful tools in detecting and quantitating individual DNA breaks in various types of experiments. An exciting area of applications is development of CRISPR/cas9 methods of genome editing. STRIDE can be applied for time-effective *in situ* validation of the efficiency and functionality of the systems used for genome editing, such as CRISPR/Cas9. Not only do these methods allow monitoring the immediate effects of the cleaving and nicking activity of the Cas9 and Cas9n proteins applied in these systems, but they could also be used to assess the level of their specificity - STRIDE can become a supplement to the existing methods for prediction and detection of off-target sites. By extending single-cell to high-throughput analysis STRIDE might prove to be a very helpful tool in speeding up the process of optimization of genome-editing systems and in their comparative analysis.

It is important to note that STRIDE may prove to be particularly useful in detecting DNA damage in cells with the mechanisms of DNA repair impaired by mutations in genes coding key repair factors, or by drugs like DDR inhibitors (such as PARP, PARG, ATM, DNA-PK inhibitors). In such cells no recruitment of some typical repair factors occur, and H2AX phosphorylation or PARylation may be weak or absent^28^. Under such conditions direct detection of DNA breaks by STRIDE could provide an unambiguous measure of DNA damage. Importantly, STRIDE bears potential to become useful in the clinic for DNA damage level assessment in patient-derived samples in the form of liquid biopsies or tissue sections. It is also worth noting that phosphorylation of H2AX histone may occur without the presence of DNA breaks^29^. In this case direct measurements of DNA damage by STRIDE may assist in avoiding overestimates of the level of damage.

As STRIDE makes it possible to detect, precisely localize inside the cell nucleus, and quantify individual DNA breaks with sensitivity that was not achievable before, we expect it to be useful in various fields of research and diagnostics, such as screening of genotoxic, anti-cancer drugs and testing their functionality and potency, basic research into mechanisms of DNA editing, DNA damage and repair and aging, in various types of medical diagnostics including infertility, and monitoring of environmental genotoxic agents. It is possible to envisage STRIDE being used in high throughput DNA damage assays for testing of new drugs. We also note that STRIDE enables recognition of the exposed DNA ends, and the technique does not have to be limited to the cell interior, thus it may prove useful in sensitive detection of circulating cell-free DNA in blood samples.

## FUNDING

This work was supported in part by Polish National Science Center (2017/27/B/NZ3/01065 to J.D., 2015/16/T/NZ3/00157 to M.K.) and U.S. National Institutes of Health grant (U01DA-040588, part of the N.I.H. 4D Nucleome Initiative) to TP.

Supplementary Data are available at NAR online.

## CONFLICT OF INTEREST

M.K., M.Z., K.S. and J.D. are the authors of a patent application covering the STRIDE methods, and co-founders of a company intoDNA.

## REFERENCES

(1) Cleaver, J. E., Feeney, L., and Revet, I. (2011) Phosphorylated H2Ax is not an unambiguous marker for DNA double-strand breaks. Cell Cycle 10, 3223–4.

(2) Rybak, P., Hoang, A., Bujnowicz, L., Bernas, T., Zarębski, M., Darzynkiewicz, Z., and Dobrucki, J. (2016) Low level phosphorylation of histone H2AX on serine 139 (γH2AX) is not associated with DNA double-strand breaks. Oncotarget 139, 1–7.

(3) Rogakou, E. P., Pilch, D. R., Orr, A. H., Ivanova, V. S., and Bonner, W. M. (1998) DNA double-stranded breaks induce histone H2AX phosphorylation on serine 139. J. Biol. Chem. 273, 5858–5868.

(4) Guo, X., Bai, Y., Zhao, M., Zhou, M., Shen, Q., Yun, C.-H., Zhang, H., Zhu, W.-G., and Wang, J. (2018) Acetylation of 53BP1 dictates the DNA double strand break repair pathway. Nucleic Acids Res. 46, 689–703.

(5) Tarsounas, M., Davies, D., and West, S. C. (2003) BRCA2-dependent and independent formation of RAD51 nuclear foci. Oncogene 22, 1115–1123.

(6) Solarczyk, K. J., Kordon, M., Berniak, K., and Dobrucki, J. W. (2016) Two stages of XRCC1 recruitment and two classes of XRCC1 foci formed in response to low level DNA damage induced by visible light, or stress triggered by heat shock. DNA Repair (Amst). 37, 12–21.

(7) Galbiati, A., Beauséjour, C., and d’Adda di Fagagna, F. (2017) A novel single-cell method provides direct evidence of persistent DNA damage in senescent cells and aged mammalian tissues. Aging Cell 16, 422–427.

(8) Ziani, S., Nagy, Z., Alekseev, S., Soutoglou, E., Egly, J. M., and Coin, F. (2014) Sequential and ordered assembly of a large DNA repair complex on undamaged chromatin. J. Cell Biol. 206, 589–598.

(9) Bekker-Jensen, S., Lukas, C., Melander, F., Bartek, J., and Lukas, J. (2005) Dynamic assembly and sustained retention of 53BP1 at the sites of DNA damage are controlled by Mdc1/NFBD1. J. Cell Biol. 170, 201–11.

(10) Kordon, M. M., Szczurek, A., Berniak, K., Szelest, O., Solarczyk, K., Tworzydło, M., Wachsmann-Hogiu, S., Vaahtokari, A., Cremer, C., Pederson, T., and Dobrucki, J. W. (2019) PML-like subnuclear bodies, containing XRCC1, juxtaposed to DNA replication-based single-strand breaks. FASEB J. 33, 2301–2313.

(11) Gorczyca, W., Bruno, S., Darzynkiewicz, R., Gong, J., and Darzynkiewicz, Z. (1992) DNA strand breaks occurring during apoptosis - their early insitu detection by the terminal deoxynucleotidyl transferase and nick translation assays and prevention by serine protease inhibitors. Int. J. Oncol. 1, 639–648.

(12) Gorczyca, W., Gong, J., and Darzynkiewicz, Z. (1993) Detection of DNA strand breaks in individual apoptic cells by the in situ terminal deoxynucleotidyltransferse and nick translation assays. Cancer Res. 53, 1945–1951.

(13) Schnell, U., Dijk, F., Sjollema, K. A., and Giepmans, B. N. G. (2012) Immunolabeling artifacts and the need for live-cell imaging. Nat. Methods 9, 152–158.

(14) Manning, C. F., Bundros, A. M., and Trimmer, J. S. (2012) Benefits and pitfalls of secondary antibodies: why choosing the right secondary is of primary importance. PLoS One 7, e38313.

(15) Crosetto, N., Mitra, A., Silva, M. J., Bienko, M., Dojer, N., Wang, Q., Karaca, E., Chiarle, R., Skrzypczak, M., Ginalski, K., Pasero, P., Rowicka, M., and Dikic, I. (2013) Nucleotide-resolution DNA double-strand break mapping by next-generation sequencing. Nat. Methods 10, 361–365.

(16) Yan, W. X., Mirzazadeh, R., Garnerone, S., Scott, D., Schneider, M. W., Kallas, T., Custodio, J., Wernersson, E., Li, Y., Gao, L., Federova, Y., Zetsche, B., Zhang, F., Bienko, M., and Crosetto, N. (2017) BLISS is a versatile and quantitative method for genome-wide profiling of DNA double-strand breaks. Nat. Commun. 8, 1–9.

(17) Biernacka, A., Zhu, Y., Skrzypczak, M., Forey, R., Pardo, B., Grzelak, M., Nde, J., Mitra, A., Kudlicki, A., Crosetto, N., Pasero, P., Rowicka, M., and Ginalski, K. (2018) i-BLESS is an ultrasensitive method for detection of DNA double-strand breaks. Commun. Biol. 1, 181.

(18) Tsai, S. Q., Zheng, Z., Nguyen, N. T., Liebers, M., Topkar, V. V, Thapar, V., Wyvekens, N., Khayter, C., Iafrate, A. J., Le, L. P., Aryee, M. J., and Joung, J. K. (2015) GUIDE-seq enables genome-wide profiling of off-target cleavage by CRISPR-Cas nucleases. Nat. Biotechnol. 33, 187–197.

(19) Lensing, S. V., Marsico, G., Hänsel-Hertsch, R., Lam, E. Y., Tannahill, D., and Balasubramanian, S. (2016) DSBCapture: In situ capture and sequencing of DNA breaks. Nat. Methods 13, 855–857.

(20) Soutoglou, E., Dorn, J. F., Sengupta, K., Jasin, M., Nussenzweig, A., Ried, T., Danuser, G., and Misteli, T. (2007) Positional stability of single double-strand breaks in mammalian cells. Nat. Cell Biol. 9, 675–82.

(21) Zarebski, M., Wiernasz, E., and Dobrucki, J. (2009) Recruitment of heterochromatin protein 1 to DNA repair sites. Cytom. Part A 75, 619–625.

(22) Solarczyk, K. J., Zarębski, M., and Dobrucki, J. W. (2012) Inducing local DNA damage by visible light to study chromatin repair. DNA Repair (Amst). 11, 996–1002.

(23) Jinek, M., Chylinski, K., Fonfara, I., Hauer, M., Doudna, J. A., and Charpentier, E. (2012) A programmable dual-RNA-guided DNA endonuclease in adaptive bacterial immunity. Science 337, 816–821.

(24) Ran, F. A., Hsu, P. D., Lin, C.-Y., Gootenberg, J. S., Konermann, S., Trevino, A. E., Scott, D. A., Inoue, A., Matoba, S., Zhang, Y., and Zhang, F. (2013) Double nicking by RNA-guided CRISPR Cas9 for enhanced genome editing specificity. Cell 154, 1380–1389.

(25) Ma, H., Naseri, A., Reyes-Gutierrez, P., Wolfe, S. A., Zhang, S., and Pederson, T. (2015) Multicolor CRISPR labeling of chromosomal loci in human cells. Proc. Natl. Acad. Sci. 112, 3002–3007.

(26) Ma, H., Tu, L. C., Naseri, A., Huisman, M., Zhang, S., Grunwald, D., and Pederson, T. (2016) CRISPR-Cas9 nuclear dynamics and target recognition in living cells. J. Cell Biol. 214, 529–537.

(27) Anglada, T., Repulles, J., Espinal, A., LaBarge, M. A., Stampfer, M. R., Genesca, A., and Martin, M. (2019) Delayed gammaH2AX foci disappearance in mammary epithelial cells from aged women reveals an age-associated DNA repair defect. Aging (Albany. NY). 11, 1510–1523.

(28) Schuhwerk, H., Bruhn, C., Siniuk, K., Min, W., Erener, S., Grigaravicius, P., Kruger, A., Ferrari, E., Zubel, T., Lazaro, D., Monajembashi, S., Kiesow, K., Kroll, T., Burkle, A., Mangerich, A., Hottiger, M., and Wang, Z.-Q. (2017) Kinetics of poly(ADP-ribosyl)ation, but not PARP1 itself, determines the cell fate in response to DNA damage in vitro and in vivo. Nucleic Acids Res. 45, 11174–11192.

(29) Tu, W.-Z., Li, B., Huang, B., Wang, Y., Liu, X.-D., Guan, H., Zhang, S.-M., Tang, Y., Rang, W.-Q., and Zhou, P.-K. (2013) γH2AX foci formation in the absence of DNA damage: mitotic H2AX phosphorylation is mediated by the DNA-PKcs/CHK2 pathway. FEBS Lett. 587, 3437–43.

